# Systemic tumor-targeting gene drive vectors proactively eliminate drug resistance in solid tumors

**DOI:** 10.1101/2025.10.04.680477

**Authors:** Scott M Leighow, Michael T Hemann, Justin R Pritchard

## Abstract

While targeted therapies have revolutionized cancer treatment, drug resistance remains a major barrier to their curative potential. We recently demonstrated biological proof-of-concept for selection gene drive circuits, a technology that overwrites disease evolution to proactively eliminate resistance in vivo, but translation requires a delivery method compatible with disseminated metastatic disease. Now, we demonstrate a clinically feasible delivery solution with novel tumor-targeting lentiviral vectors that selectively install these therapeutic circuits in tumor cells in situ. Systemic administration of these vectors demonstrated durable elimination of visible tumor burden and minimal body weight loss, validating the translational potential of a new class of genetic medicines for long-term control of resistance in cancer.

## MAIN

Precision oncology has enabled targeted therapies that demonstrate impressive safety and efficacy^1^. However, long-term survival remains low for patients with advanced solid tumors, as drug resistance limits the durability of these agents^2^. Drug combinations are proven to suppress resistance^3,4^, but safe and effective combination therapy requires multiple, tractable vulnerabilities - which most tumors lack. Large scale functional genomic screens^5,6^ have highlighted that for any individual cancer, there are very few “home run” essential targets with dramatic, single-agent efficacy like EGFR and KRAS. An effective combination would depend on more than one of these public, tractable vulnerabilities within the same tumor.

Gene therapy offers a potential solution. Rather than identifying new targets, engineering tumors with additional, synthetic vulnerabilities^7,8^ could enable tumor-specific combination treatments. For these engineered vulnerabilities to create a meaningful therapeutic window, two conditions must be met: (i) widespread genetic modification within the tumor to drive efficacy, even in the face of native tumor heterogeneity, and (ii) minimal transduction of healthy tissue to maintain safety. Towards the first requirement, we previously described the “selection gene drive” platform, which enables tumor-wide reprogramming even from low levels of initial genetic modification^9^. This approach can effectively redesign a tumor with therapeutic genes that convert otherwise inert prodrugs into local chemotherapeutics. Complete tumor elimination is enabled by bystander activity, whereby local diffusion of the activated metabolites clears the unmodified tumor fraction.

Successful translation of this approach requires a delivery vehicle that satisfies the second system requirement: tumor-specific transduction. To this end, we take advantage of recent advancements in targeted lentiviral vector (LVVs), a strategy that has been clinically validated for in vivo delivery of CAR vectors to T cells^10^. Here, we modify this new technology to create novel tumor-targeting LVVs that are programmed to selectively recognize and modify tumor cells. Systemic vector administration enables gene delivery to the tumor, without a priori knowledge of anatomic location. From there, the delivered genetic circuits reprogram and eliminate the tumor, proactively clearing resistant clones that would otherwise drive treatment failure. Rare off-target transduction is subject to counterselection, thereby reinforcing a strong therapeutic window.

First, we evaluated whether LVVs could be rational programmed to selectively target tumor cells. LVVs offer several features that make them well-suited for gene drive delivery. Their cargo capacity is compatible with gene drive circuits (7-9 kb), and system function requires the stable inheritance afforded by genomic integration^11^. Additionally, our previous gene drive studies involving ex vivo modification used lentiviral transduction, establishing system compatibility^9^.

Targeted LVVs have recently been described^12–14^. These vectors are pseudotyped with mutant VSV-G with abrogated LDLR binding but preserved fusogenic activity^15^. Tropism is programmed with scFv molecules that recognize surface markers selective to the target cell^12–14^ (Figure 1a). Targeted LVVs that engineer CAR-T cells in vivo have recently entered the clinic, underscoring the translational feasibility of this approach^10,16,17^.

**Figure 1:**
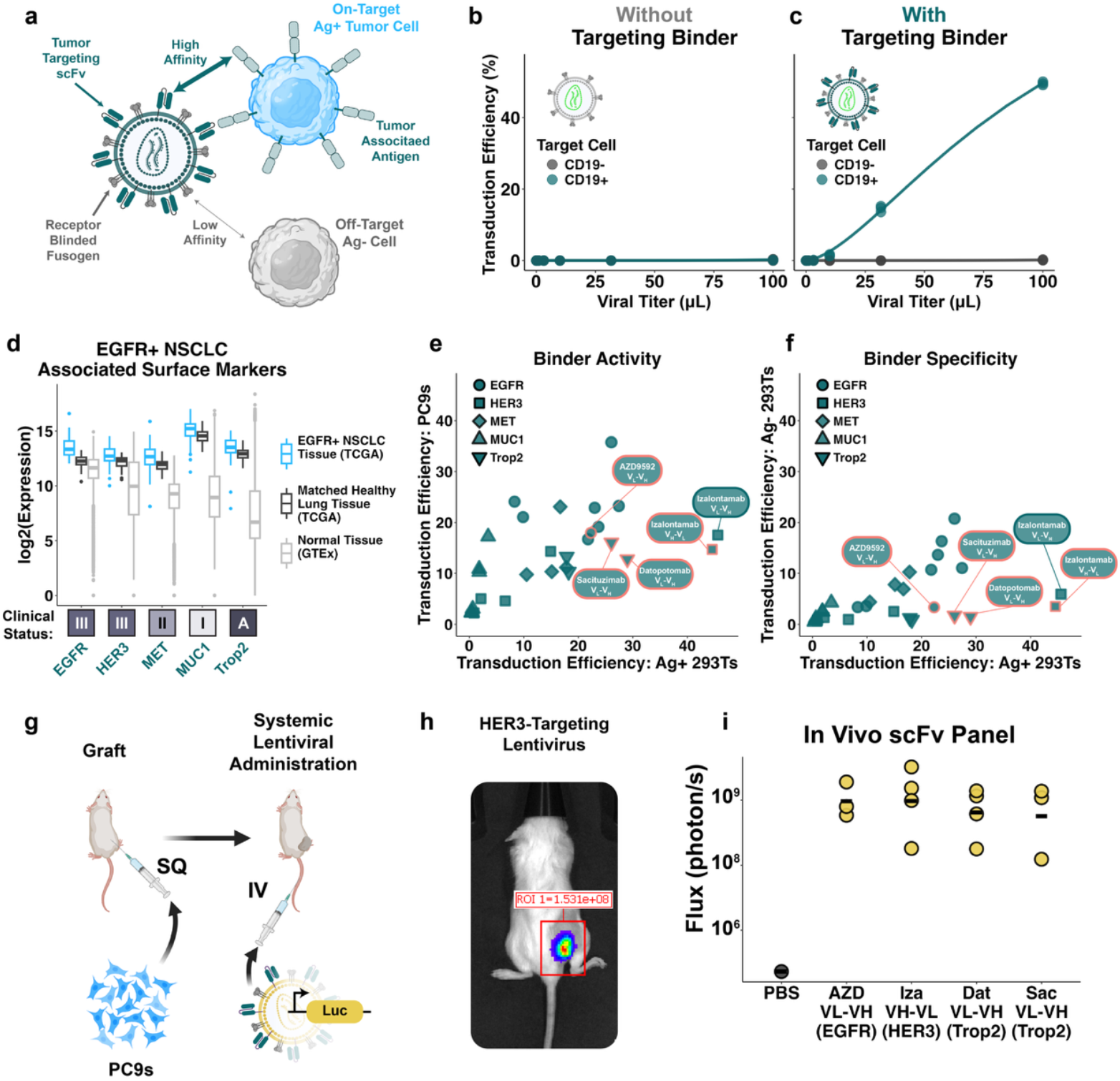
Lentiviral particles can be programmed to selectively target solid tumors. **(a)** Schematic of targeted lentiviral vector. The mutations in the receptor-blinded fusogen (VSV-G) abrogate affinity for native LDLR. Tropism is instead programmed through an scFv molecule directed against a tumor-associated antigen. **(b-c)** In vitro proof-of-concept of targeted LVVs. CD19+ (green) and wild-type, CD19- (gray) 293Ts were seeded 1:1 and treated with GFP LVVs pseudotyped with mutant VSV-G only **(b)** or mutant VSV-G and an anti-CD19 scFv **(c)**. On- and off-target transduction was measured by CD19 staining and GFP expression using flow cytometry. **(d)** Expression data for five EGFR+ NSCLC-associated targets in harmonized TCGA/GTEx transcriptomic data^28^. Boxes below indicate the clinical status of the most mature ADC for each target (phase I, II, III, or A for approved). **(e)** On-target activity of scFv candidates. Targeted LVVs pseudotyped with mutant VSV-G and candidate scFv molecules were used to transduce antigen-positive 293T cells (x-axis) and EGFR+ NSCLC PC9 cells (y-axis). **(f)** Specificity of scFv candidates. Targeted LVVs were used to transduce antigen-positive (x-axis) and wild-type, antigen-negative 293T cells (y-axis) as in **(e)**. Candidates nominated for in vivo evaluation are outlined in pink. **(g)** Schematic of in vivo systemic administration experiment. PC9 cells were subcutaneously grafted in mice; upon tumor formation, targeted LVVs carrying a luciferase transgene reporter were administered by tail vein injection. **(h)** Representative image of luciferase activity in mouse after tail vein injection of targeted LVV (izalontamab-based design targeting HER3). **(i)** Results from in vivo screen of candidate scFv molecules (n = 3-4 mice/scFv).

We first established targeted LVV functionality in our own hands, using a previously described^14^ model for targeting CD19+ cells (Figure 1b-c). To reprogram these vectors to recognize clinically relevant tumor-associated surface markers, we considered the targets of ADCs. Both LVVs^15^ and ADCs^18^ enter their target cell via receptor-mediated endocytosis, so the validated epitopes of ADCs should be strong candidates for LVV targeting. Moreover, clinically-validated ADCs confirm that these antigens provide an acceptable therapeutic window for tumor-selective delivery.

Unlike ADCs, however, gene drive therapy does not require delivery to the majority of tumor cells to achieve efficacy. As few as 10% engineered cells are sufficient to initiate tumor-wide reprogramming^9^, enabling therapeutic effect at substantially lower effective doses and reducing off-target risk. Additionally, gene drive vectors are only dosed once, and transduced cells self-eliminate during treatment, so any adverse effects are expected to diminish over time. Finally, whereas ADCs risk systemic toxicity from premature linker hydrolysis^19^, gene drive therapy relies on prodrugs that are only activated locally, further minimizing off-target effects.

To develop targeted LVVs compatible with our existing EGFR+ gene drive system, we considered the targets of ADCs that have been clinically evaluated in EGFR+ NSCLC: EGFR^20,21^, HER3^20,22^, MET^23,24^, MUC1^25^, and Trop2^26,27^. Analysis of harmonized TCGA and GTEx data^28^ confirmed the tumor-selective expression of these surface markers in EGFR+ NSCLC (Figure 1d). We identified the sequences of 14 ADCs developed against these targets (2-4 ADCs/target), to generate 28 total candidate scFv designs (both VH-VL and VL-VH orientations; see Supplementary Table 1).

To screen for the on-target activity of these molecules, we measured infection efficiency in PC9 cells (EGFR+ NSCLC) as well as 293T cells engineered to overexpress the cognate antigen (Figure 1e). Similarly, to evaluate their specificity, we compared activity in on-target, antigen-positive 293Ts to isogenic, wild-type cells (Figure 1f). From these assays, we selected four candidates for further in vivo evaluation based on their strong activity, high specificity, and representation across distinct targets.

The clinical progress of intravenously administered CD3-targeting LVVs for in vivo CAR-T therapy^10^ suggested that systemic delivery may be similarly feasible for tumor-targeting LVVs. Systemic administration is uniquely positioned for impact in a disseminated disease like advanced cancer with (perhaps undetected) metastatic lesions. Furthermore, intravenous delivery is simpler to implement than intranodal injection, a barrier that has limited the clinical adoption of oncolytic viruses like Imlygic^29^.

To assess the feasibility of using systemic targeted LVVs to modify tumor cells in vivo, we subcutaneously grafted PC9 cells in the flanks of NSG mice. Upon tumor formation, luciferase LVVs pseudotyped with our top-performing scFv molecules were administered by tail vein injection (Figure 1g). In vivo imaging confirmed luciferase delivery to the tumor (Figure 1h), and there was no significant difference in luminescence signal strength across the four candidate scFv molecules (Figure 1i; Supplementary Figure 1). We selected the izalontamab-based design targeting HER3 for further studies based on its high on-to off-target transduction ratio in vitro and the strong clinical data supporting HER3 as a therapeutic target^30^.

Having validated an appropriate tumor-targeting delivery strategy, we next sought to optimize the gene drive circuitry itself for efficient delivery. Selection gene drives carry two genetic “switches” (Figure 2a) that operate in sequential phases^9^. In the first phase, Switch 1 functions as an inducible drug target analog; a clinically-validated dimerizer activates Switch 1 to transiently rescue engineered cells from targeted therapy action. The result is a tumor reprogrammed to be majority-engineered cells. In the second phase, Switch 2 - the therapeutic payload - converts inert prodrugs into cytotoxic metabolites that diffuse locally to eliminate both engineered and residual, unmodified cells. Together, this two-step approach proactively combats resistance, even from limited initial transduction.

**Figure 2:**
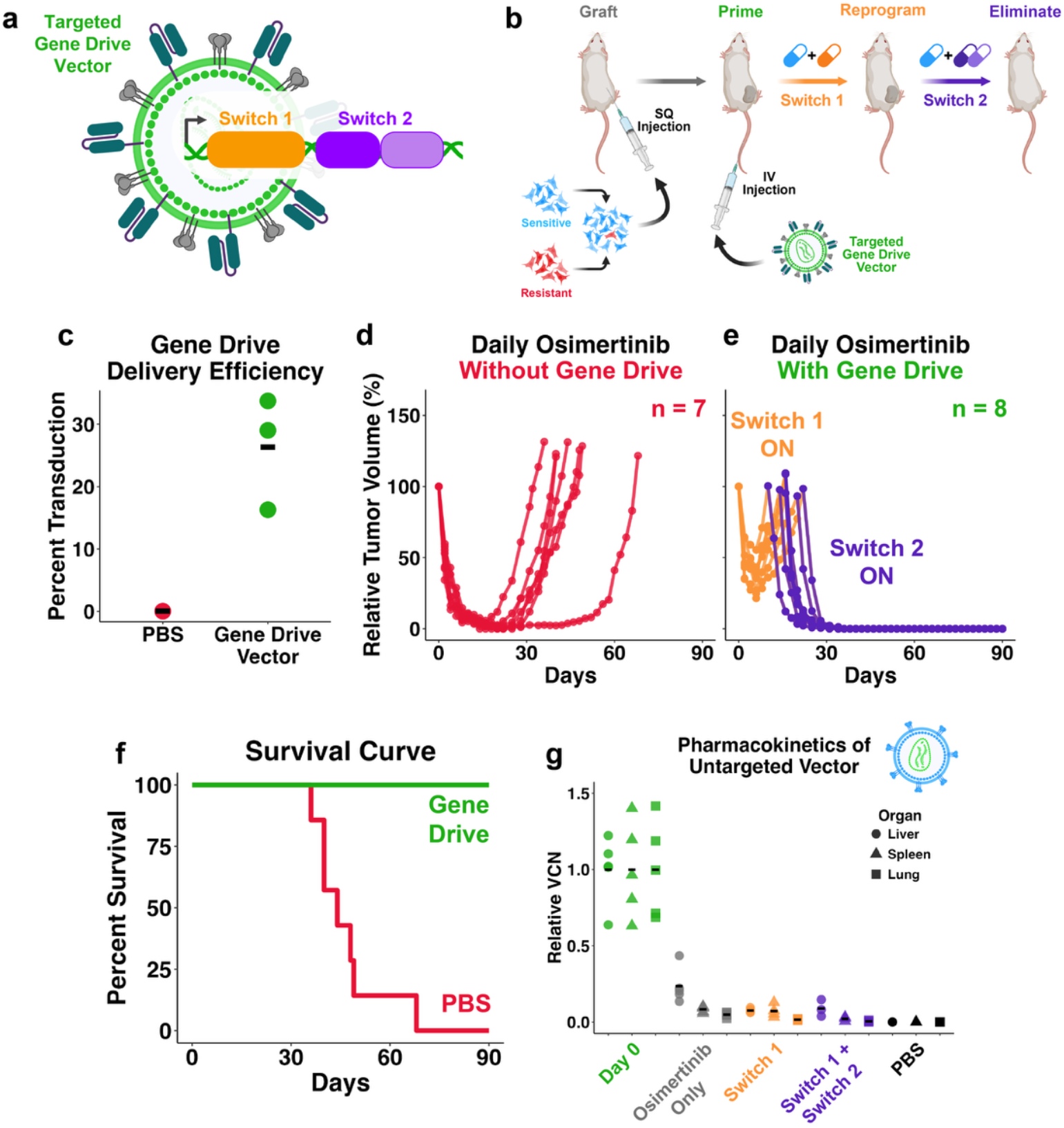
Systemic administration of optimized gene drive circuits achieves long-term disease control in vivo. **(a)** Schematic of tumor-targeting gene drive vectors. The genetic circuit composed of Switch 1 and Switch 2 elements is delivered via targeted LVV. **(b)** Schematic of in vivo gene drive vector experiment. PC9 cells (with spiked-in pre-existing resistance) were subcutaneously grafted in the flanks of mice. Upon tumor formation, tumor-targeting gene drive vectors were administered by tail vein injection. Osimertinib and the Switch 1 dimerizer molecule were administered to reprogram the tumor. Then, osimertinib and Switch 2 prodrugs were administered in combination to eliminate the tumor. **(c)** On-target dual-payload gene drive transduction efficiency, as measured by qPCR in day 0 tumors. **(d-e)** In vivo tumor dynamics. Volumes for tumors in mice primed with systemic administration of targeted dual-payload gene drive vectors **(e)** or PBS **(d)** are shown. In **(d)** line color indicates treatment with osimertinib and dimerizer (Switch 1; orange) or osimertinib and prodrugs (Switch 2; purple). **(f)** Survival curve for systemic gene drive experiment. **(g)** Pharmacokinetics of off-target transduction in liver, spleen, and lung. Untargeted gene drive vectors (wild-type VSV-G pseudotyping) were tail vein injected in tumor free mice. Mice were treated with osimertinib for three weeks (gray), osimertinib and dimerizer for three weeks (Switch 1; orange), or osimertinib and dimerizer for three weeks followed by osimertinib and prodrugs for six weeks (Switch 1+2; purple). VCN was measured by qPCR and normalized to mean day 0 values.

To adapt these circuits for systemic in vivo delivery, we redesigned them to reduce vector size while enhancing potency. First, we minimized the Switch 1 gene to decrease cargo size and consequently improve delivery efficiency. Then, we developed a dual-payload Switch 2 system by combining two previously validated^9^ suicide genes: cytosine deaminase (CyD) and the nitroreductase NfsA.This design strengthens bystander activity, thereby lowering the theoretical delivery threshold, since fewer engineered cells are needed to achieve a tumor-wide effect. Full design and validation details are provided in Supplementary Note 1 and Supplementary Figure 2.

Having individually optimized the vector and circuits for delivery, we next assembled them into a completed prototype. First, we tested the principle of targeted gene drive vectors in vitro. To test robustness under heterogeneous antigen expression, we returned to the CD19-targeting LVV model. This provided a tractable system for controlled variation in target expression, reflecting the heterogeneity observed in real tumors^31^. Mixed PC9 populations with 25% CD19+ cells were infected with CD19-targeting gene drive vectors, and subpopulation dynamics were monitored by flow cytometry. Despite transducing a minority of cells, the gene drive system eliminated pre-existing resistance in vitro (Supplementary Figure 3a-c).

We next evaluated whether tumor-targeted gene drive vectors could achieve durable disease control in vivo. Dual-payload (CyD/NfsA) circuits were packaged in HER3-targeting LVVs and then tail-vein injected in mice bearing PC9 xenografts (Figure 2b). Analysis of day 0 tumors indicated a delivery efficiency of ∼25% (mean VCN = 0.26, Figure 2c). Mice were then treated daily with the standard-of-care EGFR inhibitor osimertinib, alongside the dimerizer molecule to engage Switch 1. Switch 2 prodrugs were administered once the tumors returned to >90% of their initial volume. In mice that did not receive the gene drive vector, tumors initially regressed and then relapsed within 30-60 days, indicating they had become refractory to osimertinib (Figure 2d). However, in gene drive-primed mice, ∼20 days of Switch 1 activity successfully reprogrammed tumors into a sensitized state. Subsequent prodrug treatment resulted in deep responses by day 30, with no disease recurrence by day 90 (Figure 2e) and significantly extended survival (Figure 2f).

We similarly tested a single-payload circuit design (CyD only) in vivo. As expected, the delivery efficiency of this smaller cargo was higher than the dual-payload system (mean VCN = 0.52, Supplementary Figure 3d), but tumor regression was slower. Nonetheless, both designs prevented treatment failure over 90 days of treatment and monitoring (Supplementary Figure 3e). Additionally, neither version showed meaningful weight loss during the Switch 2 phase of treatment, suggesting no overt toxicity associated with Switch 2 function (Supplementary Figure 3f).

Next, we sought to evaluate the safety of systemically administered gene drive vectors. Their design offers several potential safety advantages over ADCs, but an important concern is whether Switch 1 activity could promote transformation in off-target cells. However, we hypothesized that the fate of gene drive transduction may differ in on- and off-target cells. While Switch 1 drives selection in malignant cells wired for strong mitogenic signaling, its activity may in fact counterselect in healthy tissue via oncogene-induced senescence^32^. Furthermore, Switch 2 may provide an additional failsafe by eliminating rare, residual off-target gene drive cells during prodrug treatment. In this way, the native function of both switches may cause gene drive circuits to be self-eliminating for off-target delivery, thereby dramatically expanding the therapeutic window in vivo.

Due to the selectivity of our HER3 vector, its off-target delivery is challenging to study in vivo. Instead, we opted to study off-target dynamics in an extremely conservative “worst case scenario” of off-target delivery: intravenous administration of a non-targeted LVV. We packaged the gene drive circuit in LVVs pseudotyped with wild-type VSV-G and administered them intravenously to tumor-free mice. The mice were then treated with either osimertinib alone, osimertinib plus dimerizer (Switch 1 activation), or osimertinib plus dimerizer followed by prodrugs (Switch 2 activation). qPCR analysis showed VCN declined across the liver, spleen, and lungs in all groups (Figure 2g). Notably, Switch 1 activation further reduced VCN in the liver and lungs, consistent with counterselection. Importantly, no increase in VCN was detected after nine weeks of treatment (Switch 1 + Switch 2 arm) in any tissue - rather these findings indicate substantial depletion of off-target transduced cells, supporting the tolerability of intravenous gene drive vector administration.

These findings establish systemic, tumor-targeted gene drive vectors as a safe and effective strategy for eliminating tumors otherwise destined for relapse. As the first full prototype of a gene drive therapy, this work defines a therapeutic framework for strategies that overwrite disease evolution and drive durable responses.

## METHODS

### Plasmid construction

Construction of the original EGFR gene drive circuit (pLVX-S1vEGFRosi-S2vCyD) was described previously^9^. PCR-based cloning was used to make the described changes, including removal of one of the tandem DmrB domains, cloning of the promoter-CDS sequence into the minimized pSL vector, and insertion of the NfsA gene. Luciferase and surface marker genes used in this study (CD19, EGFR, HER3, MET, MUC1, and Trop2) were synthesized by Genscript and cloned into the pSL vector along with a 3’ 2A-Puro sequence. To generate the panel of scFv sequences, ADC V_H_ and V_L_ amino acid sequences were obtained from IMGT or SAbDab, codon optimized for human expression, and cloned into the pJRH-scFv entry plasmid in both possible orientations (V_H_-V_L_ and V_L_-V_H_) by Genscript. Proper assembly was confirmed by whole plasmid sequencing.

### Lentiviral production

pLVX or pSL constructs were co-transfected with psPAX2 and pMD2.G in lentiX cells (Takara) using LT-1 transfection reagent (MirusBio). For targeted lentiviral production, pMD2.G was replaced with pMD2.G-K47Q/R354Q^13^ and the appropriate scFv plasmid was added. The viral supernatant was collected at 48 hours, passed through a 0.45 μm filter, and used immediately or stored at -80C. For in vitro studies, pure populations were generated under puro selection at the lowest dose sufficient to eliminate mock-infected cells. For in vivo studies, clarified viral supernatant was concentrated using sterilized 70 mL 100 kDa centricon filters (Sigma) spun at 4k g for one hour. Viral titer was measured by treating reporter GFP_1-10_ PC9 cells (gene drive vectors express a short GFP_11_ fragment) with titered virus for 24 hours. GFP expression was measured by flow cytometry after another 24 hours.

### In vitro CD19-targeting LVV validation

CD19-targeting LVVs carrying a GFP reporter gene were generated as described above using a previously validated CD19 scFv^14^. 293T cells were transduced to express CD19. CD19+ and wild-type, CD19-293Ts were mixed 1:1 and seeded in 96-well plates. 24 hours after seeding, CD19-targeting GFP lentivirus was added at diluting titers in duplicate, then removed after another 24 hours. The following day, cells were resuspended and stained with an anti-CD19/APC antibody and analyzed by flow cytometry. APC signal was used to distinguish on-target CD19+ cells, and GFP signal was used to distinguish transduced cells.

### Clinical transcriptomic data analysis

Harmonized TCGA/GTEx data was accessed via the UCSC Xena browser. TCGA data was filtered for primary lung cancer samples (and matched healthy tissue) with exon 19 deletions or L858R mutations. Gene expression data was downloaded for EGFR, ERBB3 (HER3), MET, TACSTD2 (Trop2), and MUC1 and visualized in R.

### In vitro scFv screen

LVVs carrying a GFP reporter gene were pseudotyped with mutant VSV-G and the appropriate scFv molecule. 293Ts were transduced to overexpress one of five EGFR+ NSCLC-associated markers. PC9, marker-positive 293Ts and wild-type 293Ts were seeded separately in 96 well plates. Targeted LVVs were added to the PC9, wild-type 293T, and appropriate marker-positive 293T wells in duplicate. Virus was removed after 24 hours of treatment. The following day, cells were resuspended and transduction efficiency was measured by flow cytometry.

### Mouse experiments

All animal studies were approved by the Pennsylvania State University’s Institutional Animal Care and Use Committee. Mice were housed at ambient room temperature in a humidity-controlled animal facility. Mice had free access to food and water. Mice were maintained on a 12:12 hr light:dark cycle.

### In vivo scFv screen and bioluminescence imaging

Four top-performing hits from the in vitro screen were nominated for in vivo studies: the V_L_-V_H_ AZD9592-based EGFR-targeting molecule, the V_L_-V_H_ sacituzimab-based Trop2-targeting molecule, the V_L_-V_H_ datopotomab-based Trop2-targeting molecule, and the V_H_-V_L_ izalontamab-based HER3-targeting molecule. NSG mice (Jackson Labs) were subcutaneously grated with PC9 cells and assigned to groups to minimize differences in tumor volume mean and standard deviation. 200 µL of targeted luciferase LVVs or PBS were tail-vein injected every day for three days. 48 hours after the last injection, luminescence activity was visualized and quantified with an IVIS imager.

### In vitro Switch 1 response assay

PC9 cells were transduced with alternative gene drive circuit designs (i.e. single vs tandem S1vEGFR_osi_ dimerization domains, pLVX vs pSL lentiviral backbones) and purified by cell sorting as previously described^9^. Cells were seeded in 96-well plates and treated with osimeritinib (1 nM to 10 µM) and dimerizer (10 nM) in triplicate. Cell viability was measured after 72 hours using CellTiter-Glo 2.0 (Promega) and normalized to untreated controls.

### In vitro Switch 2 dose response

PC9 cells were transduced with complete gene drive circuits carrying either single-(CyD) or dual-payloads (CyD+NfsA) and purified by cell sorting as previously described^9^. Cells were seeded in 96-well plates and, after 24 hours, treated with 5-FC (1-10 mM) or CB1954 (31.6-316 µM). Cell viability was measured after 72 hours of drug treatment using CellTiter-Glo 2.0 (Promega) and normalized to untreated controls.

### In vitro Switch 2 bystander effect

Wild-type and gene drive (single- or dual-payload) PC9 cells were mixed for a range of gene drive ratios (0%, 12.5%, 25%, 50%, and 100%) and seeded in 96-well plates. After 24 hours, cells were treated with 10 mM 5-FC, 316 µM CB1954, or both. Cell viability was measured after 48 hours of drug treatment using CellTiter-Glo 2.0 (Promega) and normalized to untreated controls.

### In vivo Switch 2 bystander effect

PC9 cells transduced with either the single- (CyD) or dual-payload (CyD+NfsA) gene drive systems were mixed with wild-type PC9 cells at 1:1 ratio to reflect partial tumor remodeling. These mixed populations were subcutaneously grafted in the flanks of mice. Upon tumor establishment, mice were i.p. injected daily with either 500 mg/kg 5-FC (prepared in sterile PBS), 20 mg/kg CB1954 (in 2% Tween), or both. Mice also received daily 25 mg/kg osimertinib (oral gavage; formulated in NMP/PEG) to match the dosing regimen associated with the Switch 2 phase of treatment. Tumor volumes were regularly measured by calipers.

### In vitro targeted gene drive vector proof-of-concept

Pooled populations of PC9 cells were generated by mixing 74.5% wild-type, CD19-cells, 25% CD19+ (targetable) cells, and 0.5% EGFR L858R/C797S, mCherry+ cells, and seeded into 24-well plates. After 24 hours, cells were treated with gene drive vectors pseudotyped with the CD19-targeting scFv. Virus was removed after another 24 hours, and the populations were treated with the osimertinib/dimerizer and osimertinib/prodrug regimen as described previously^9^. Subpopulation dynamics were regularly monitored by flow cytometry.

### In vivo targeted gene drive therapy

For the on-target study, single- and dual-payload gene drive circuits were packaged in HER3-targeted LVVs. Mixed populations of sensitive and resistant (.01%) PC9 cells were generated ex vivo and subcutaneously grafted in NSG mice. Upon tumor establishment, mice were assigned to one of three groups (single-, dual-payload gene drive, or PBS) to minimize differences in tumor volume mean and standard deviation. Then, mice were tail vein injected with 200 µL of the appropriate targeted vector for five days. After gene drive conditioning, a subset of three mice from each arm were euthanized and their tumors harvested for further analysis (see qPCR methods). The remaining mice received 25 mg/kg osimertinib (oral gavage; prepared in NMP/PEG) and 1 mg/kg dimerizer (i.p. injection; prepared in 2% Tween) daily, until tumors returned to >90% their original volume. At this point, mice received osimertinib and 5-FC (single-payload arm; 5-FC dosed at 500 mg/kg) or osimertinib, 5-FC, and CB1954 (dual-payload and control PBS arms; CB1954 dosed at 20 mg/kg). Mice also continued to receive dimerizer treatment for the first three weeks of the Switch 2 prodrug phase, because prior studies demonstrated that such an overlap in treatment improved bystander function^9^. Tumor volumes were monitored regularly with calipers.

For the off-target study, the single-payload gene drive circuit was packaged in untargeted, wild-type VSV-G pseudotyped LVVs. NSG mice were tail vein injected with 200 uL of the untargeted gene drive vector (or PBS) for five days and randomized to one of several groups: a day 0 arm to establish baseline VCN values; an arm receiving three weeks of osimertinib (25 mg/kg); an arm receiving three weeks of osimertinib plus dimerizer (1 mg/kg); and an arm receiving three weeks of osimertinib plus dimerizer followed by six weeks of osimertinib plus 5-FC (500 mg/kg). At each terminal timepoint, liver, spleen, and lung tissue were collected.

### On- and off-target vector copy number (VCN) qPCR measurement

Tumor and organ samples were harvested at terminal timepoints, pulverized in liquid nitrogen, and stored as powder in -20C. gDNA was later extracted (Qiagen) and gene drive copy number was measured by qPCR (100 ng DNA/well in duplicate). Because spiked-in resistant PC9 cells were also transduced by lentivirus (ex vivo), common lentiviral qPCR probes could not be used to specifically measure gene drive provirus. Instead, custom probes were designed against CyD. Additionally, for tumor sample specifically, human RNase P probes (ThermoFisher) were used as an endogenous control to determine human gDNA content in each well.

## SUPPLEMENTARY TEXT

### Supplementary Note 1

To improve delivery efficiency, we sought to minimize the cargo size of our gene drive circuits without compromising function. The initial Switch 1 design, derived from a base Clontech vector, utilized dual-tandem dimerization (DmrB) domains. We tested whether a single DmrB domain could provide equivalent functionality (Supplementary Figure 2a). In fact, the single-domain design showed modestly improved performance, with lower basal activity and higher responsiveness upon dimerizer treatment compared to the dual-tandem system (Supplementary Figure 2b).

Next, we examined whether a smaller lentiviral backbone could further reduce vector size. The standard pLVX backbone has a total length of 7.9 kb LTR-LTR, whereas the alternative pSL backbone is 6.8 kb. We therefore cloned our promoter-CDS cassette into the pSL backbone (Supplementary Figure 2c) and confirmed that Switch 1 function was fully preserved (Supplementary Figure 2d).

Finally, we investigated whether incorporating two distinct Switch 2 payload genes could enhance therapeutic performance. The rationale was twofold: first, stronger bystander activity would reduce the delivery threshold, since fewer engineered cells would be needed to achieve tumor-wide reprogramming; second, using two independent prodrug-activating enzymes would minimize the risk of cross-resistance. In addition to the cytosine deaminase (CyD) gene, which we had validated previously, we selected the nitroreductase NfsA, an enzyme that activates the prodrug CB1954. NfsA had shown both sensitivity to CB1954 and robust bystander activity in our prior work^9^. We cloned both genes into the gene drive vector, which increased the cargo size from 6.8 kb to 7.6 kb (Supplementary Figure 2e). As expected, the larger cassette resulted in a modest reduction in infection efficiency (Supplementary Figure 2f). However, the addition of NfsA did not compromise CyD function: sensitivity to 5-FC was unchanged between the single-payload (CyD) and dual-payload (CyD/NfsA) systems (Supplementary Figure 2g). Moreover, the dual system demonstrated sensitivity to CB1954, validating the functionality of NfsA (Supplementary Figure 2h).

We next evaluated the bystander activity of the dual-payload system. CyD-mediated 5-FC bystander activity was equivalent between single- and dual-payload designs, while NfsA conferred robust CB1954 bystander activity on its own. Most strikingly, when both prodrugs were combined (5-FC + CB1954), the dual-payload system achieved nearly complete elimination of mixed cell populations even when only 25% of cells were engineered (Supplementary Figure 2f). Finally, we confirmed the in vivo efficacy of the dual-payload design in a mixed tumor model. Xenografts seeded with a 1:1 mixture of wild-type and gene drive cells exhibited strong Switch 2 activity and tumor regression following prodrug treatment (Supplementary Figure 2g).

## SUPPLEMENTARY FIGURES

**Supplementary Figure 1:**
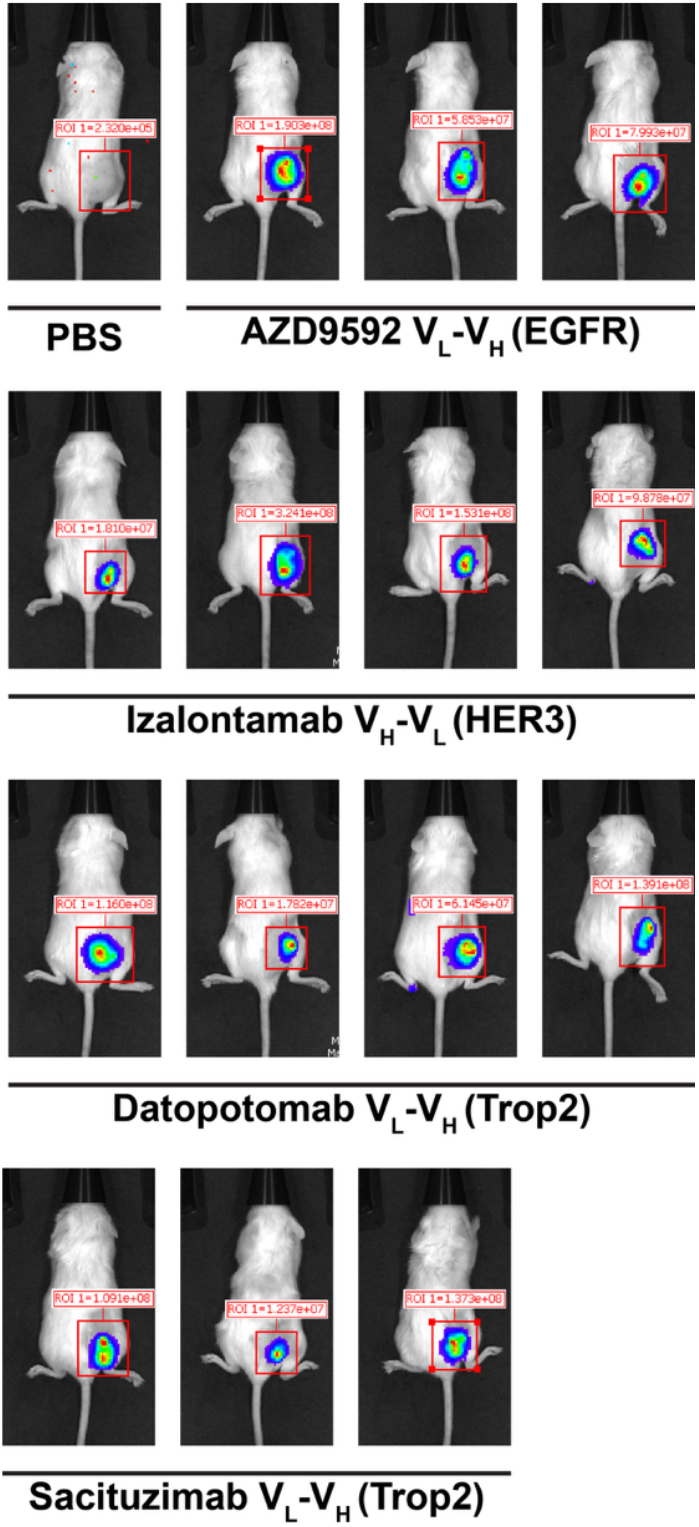
Bioluminescence imaging for in vivo scFv screen. PC9 cells were subcutaneously grafted in the left flanks of mice. Targeted luciferase LVVs pseudotyped with the noted scFv molecules were administered by tail vein injection.

**Supplementary Figure 2:**
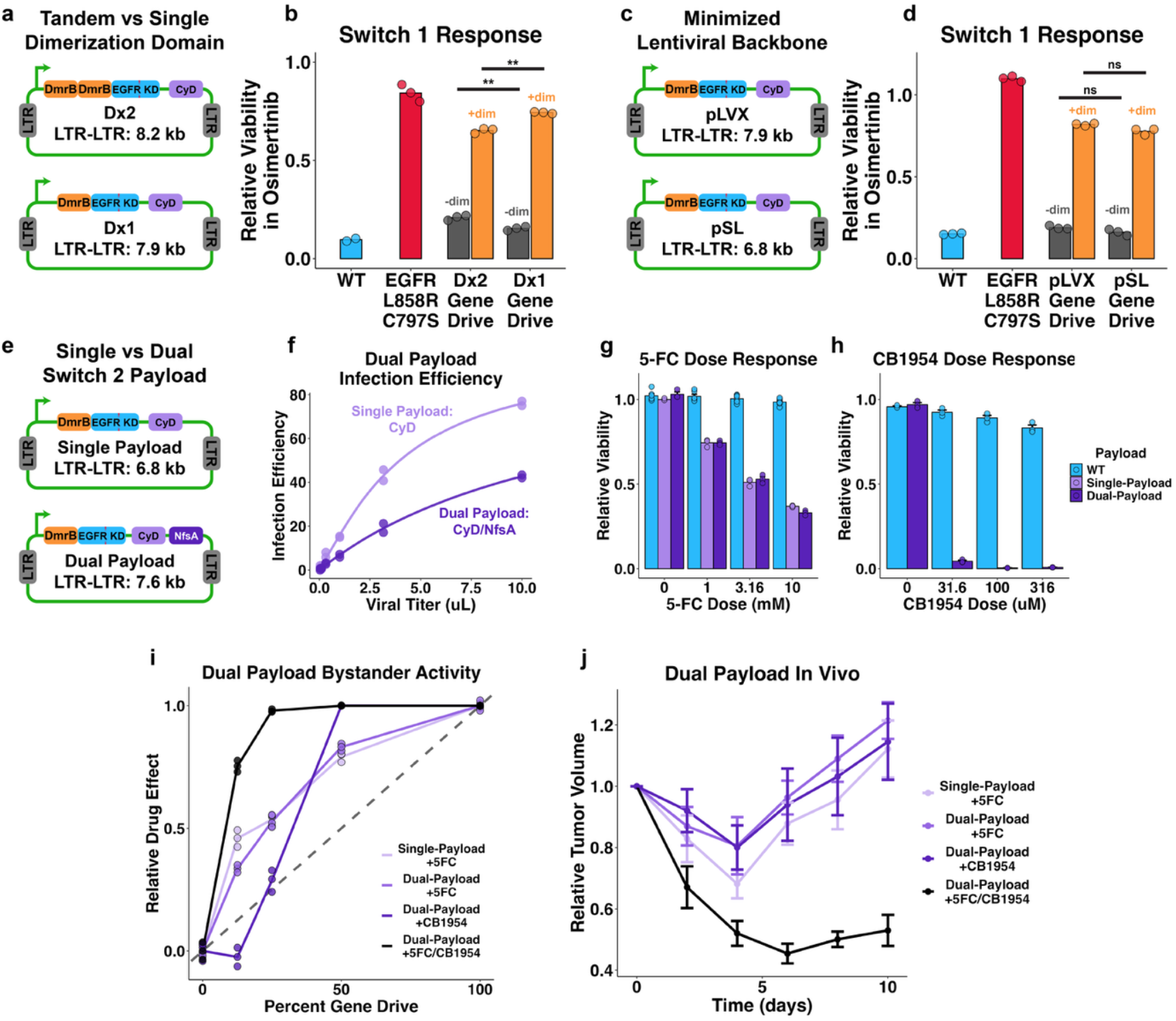
Optimization of gene drive circuits for delivery. **(a)** Schematic plasmid maps for the tandem- (original) and single-dimerization domain Switch 1 designs. **(b)** Switch 1 activity for the tandem- (Dx2) and single-dimerization (Dx1) domain designs. PC9 cells were transduced with either circuit and treated with osimertinib with (orange) or without (gray) dimerizer. Wild-type (blue) and resistant EGFR L858R/C797S (red) serve as controls. **(c)** Schematic plasmid maps for gene drive circuits in pLVX and pSL lentiviral plasmid vectors. **(d)** Switch 1 activity for the pLVX and pSL versions of the gene drive system, as in **(b)**. **(e)** Schematic plasmid maps for the single- (CyD only) and dual-payload (CyD+NfsA) gene drive circuits. **(f)** Transduction efficiency of the single- and dual-payload circuits. Reporter GFP_1-10_ PC9 cells were infected with either design (carrying a small GFP_11_ fragment) for a range of viral titers. Transduction efficiency was measured by flow cytometry. **(g-h)** Dose response for the prodrugs 5-FC **(g)** and CB1954 **(h)**. Data shown for wild-type (blue), single-payload (light purple), or dual-payload (dark purple) PC9 cells. **(i)** Bystander curves for single- and dual-payload systems. Mixed PC9 populations were seeded at various percentages of gene drive cells and treated with prodrug. In the absence of bystander function, the relative drug effect should be proportional to the fraction of gene drive cells expressing the payload suicide gene. Observed drug effect greater than the line of unity indicates bystander function. Data shown for single-payload cells treated with 5-FC (light purple), dual-payload cells treated with 5-FC (medium purple), dual-payload cells treated with CB1954 (dark purple), and dual-payload cells treated with both 5-FC and CB1954 (black). **(j)** In vivo tumor dynamics in response to prodrug treatment. Wild-type and gene drive (single- or dual-payload) PC9 cells were mixed 1:1 ex vivo and grafted in mice. Upon tumor formation, mice were treated with 5-FC and/or CB1954. Data shown for single-payload mice treated with 5-FC (light purple), dual-payload mice treated with 5-FC (medium purple), dual-payload mice treated with CB1954 (dark purple), and dual-payload mice treated with 5-FC and CB1954 (black).

**Supplementary Figure 3:**
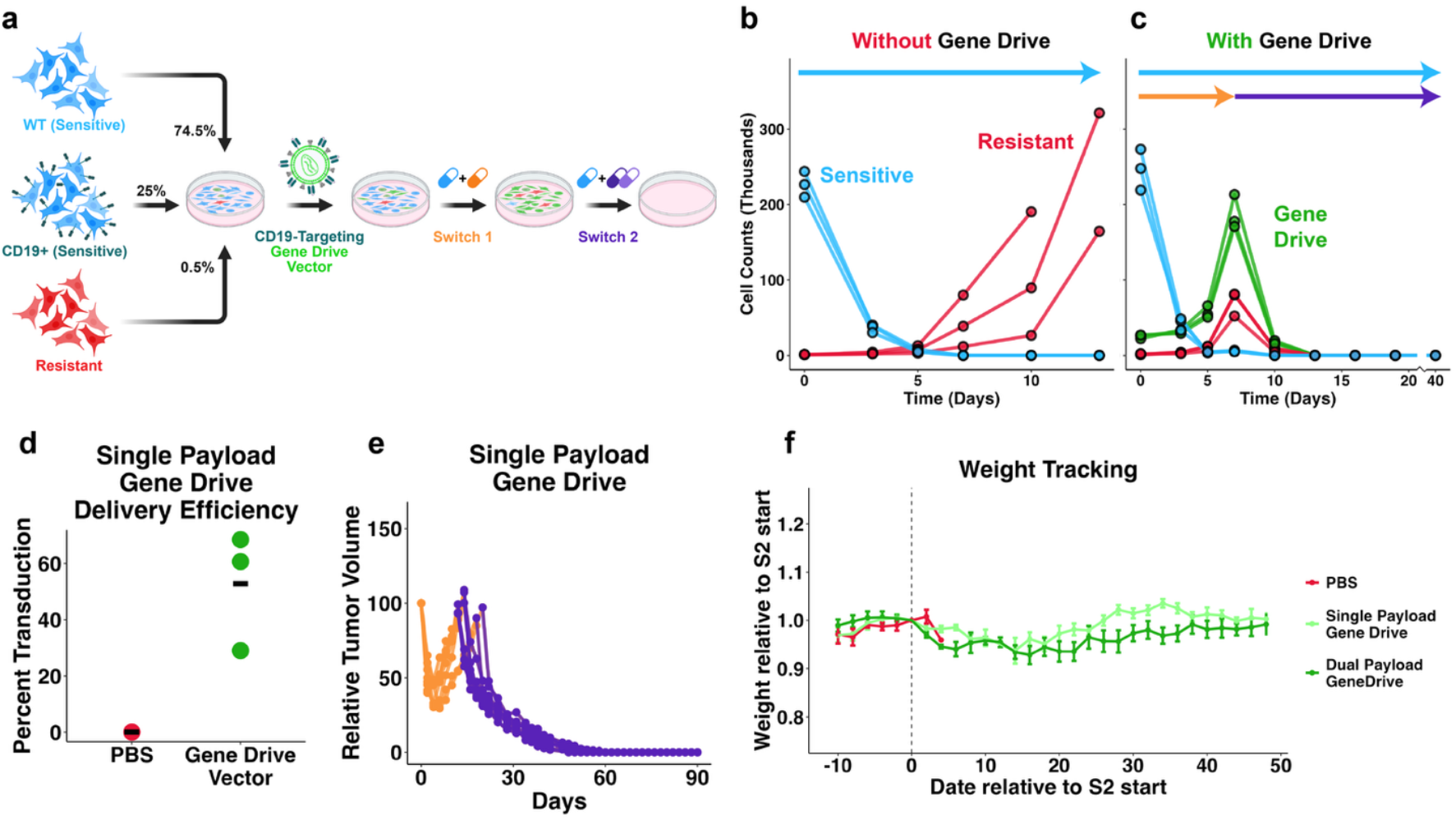
Integration of gene drive circuits with targeted LVVs. **(a)** Experimental schematic of in vitro targeted gene drive vector experiment. Wild-type, CD19+ (targetable), and resistant (EGFR L858R/C797S, mCherry+) PC9 cells were pooled and seeded. The mixed populations were briefly treated with CD19-targeting gene drive LVVs. After removal of virus, cells were treated with osimertinib and dimerizer. Upon outgrowth of gene drive cells, the mixed population was treated with osimertinib and 5-FC/CB1954. **(b-c)** Functionality of targeted gene drive LVVs in vitro. Populations treated with **(c)** and without **(b)** targeted gene drive vectors are shown. Flow cytometry was used to monitor sensitive (blue), resistant (red), and gene drive (green) subpopulation dynamics. Blue, orange, and purple arrows indicate osimertinib, dimerizer, and 5-FC treatment, respectively. **(d)** On-target single-payload gene drive transduction efficiency, as measured by qPCR in day 0 tumors, as in Figure 2c. **(e)** In vivo tumor dynamics for mice primed with systemic administration of targeted single-payload drive vectors, as in Figure 2e. Line color indicates treatment with osimertinib and dimerizer (Switch 1; orange) or osimertinib and prodrugs (Switch 2; purple). **(f)** Monitoring of mouse weight in in vivo targeted gene drive vector experiment. Time is translated on the x-axis relative to the start of Switch 2 (prodrug) treatment for each animal. Values are normalized to the weight at the start of Switch 2 treatment. Data are shown for mice primed with control PBS (red), single- (light green) and dual-payload (dark green). Note that control PBS mice were generally euthanized 2-4 days after the start of Switch 2 treatment (when tumors reached >90% original volume), due to continued disease progression.

